# Morphological and genetic diversity of maize landraces along an altitudinal gradient in the Southern Andes

**DOI:** 10.1101/2022.07.01.498464

**Authors:** Juan G. Rivas, Angela V. Gutierrez, Raquel A. Defacio, Jorge Schimpf, Ana L. Vicario, H. Esteban Hopp, Norma B. Paniego, Veronica V. Lia

## Abstract

Maize (*Zea mays* ssp. *mays)* is a major cereal crop worldwide and is traditionally or commercially cultivated almost all over the Americas. The northwestern region of Argentina (NWA) constitutes one of the main diversity hotspots of the Southern Andes, with contrasting landscapes and a large number of landraces. Despite the extensive collections performed by the “Banco Activo de Germoplasma INTA Pergamino, Argentina” (BAP), most of them have not been characterized yet. Here we report the morphological and molecular evaluation of 30 accessions collected from NWA, along an altitudinal gradient between 1120 and 2950 meters above sea level (masl). Assessment of morphological variation in a common garden allowed the discrimination of two groups, which differed mainly in endosperm type and overall plant size. Although the groups retrieved by the molecular analyses were not consistent with morphological clusters, they showed a clear pattern of altitudinal structuring. Affinities among accessions were not in accordance with racial assignments. Overall, our results revealed that there are two maize gene pools co-existing in NWA, probably resulting from various waves of maize introduction in pre-Columbian times as well as from the adoption of modern varieties by local farmers.

In conclusion, the NWA maize landraces preserved at the BAP possess high morphological and molecular variability. Our results highlight their potential as a source of diversity for increasing the genetic basis of breeding programs and provide useful information to guide future sampling and conservation efforts.

## Introduction

Genetic erosion is the main problem associated with the selection and improvement of agronomically important species. Despite the progress achieved in terms of productivity, modern breeding takes advantage of only a small fraction of the available variability, restricting the response of crops to pests, diseases and environmental changes [1–3]. The use of landraces or related wild species provides the opportunity to counteract this process, widening the narrow genetic base of elite germplasm.

The recent revalorization of landraces as sources of diversity and beneficial alleles has been expressed through different initiatives aimed at providing an extensive characterization of germplasm bank collections. This characterization involves molecular and phenotypic aspects of maize and other crops [e.g., https://seedsofdiscovery.org/; www.amaizing.fr;, 4]. The high genetic diversity of maize landraces in the Americas has been thoroughly documented. [e.g. 5–11]. So far, however, most studies have adopted a macro-regional approach, where large geographic areas are represented by a small number of accessions and each accession is represented by a small number of individuals or sample pools. The information provided by this type of strategy is extremely valuable in characterizing variability, but it is insufficient for other genetic population analyses, especially those linked to environmental variables and local adaptations. Moreover, the limited number of studies aiming to examine the relationship among phenotype, genotype and environment through the combination of molecular and morphological data may be accounted for by the difficulty in seed germination and growth of landraces outside their native range.

The northwestern region of Argentina (NWA) is the southernmost distribution limit of the Andean maize landraces. This region is characterized by a remarkable topographic variability, as it comprises six phytogeographic provinces located within a relatively limited area (i.e, Yungas, Chaco, Puna, Pre-Puna, Monte and High Andean) [12]. Two contrasting examples of its great environmental diversity are the subtropical forests of the Yungas (distributed between 400 and 3000 masl and with annual mean precipitation increasing between 600 and 3000 mm with altitude), and the ridges and valleys of the Sub-Andean mountains (precipitation below 200 mm/ year, mainly concentrated in the summer months)[12].

More than 50% of the *ca*. 56 maize landraces described for northern Argentina are native to NWA, making it one of the main diversity hotspots of the Southern Andes [13,14]. In accordance, the most ancient records of maize from the American Southern Cone correspond to NWA and date to 3500 years BP [15]. Currently, the native germplasm is traditionally cultivated between 400 and 3600 masl, thus being exposed to a wide range of thermal amplitudes and rainfall regimes [13]. So far, only a limited number of maize landraces from NWA have been characterized either molecularly and/or cytogenetically [16,17]. Although most of these races are associated with the Andean Complex, as defined by Mc. Clintock et al. [18], the presence of germplasm from other origins (e.g. popcorn and tropical maize) was also detected [16,17].

The present scenario of increasing food demand and climate change highlights the need for materials capable of growing under extreme conditions, such as the NWA landraces. Therefore, the characterization of the accessions in germplasm banks is important in tackling these challenges. Today, the “Banco Activo de Germoplasma de maíz in EEA-INTA Pergamino, Argentina” (BAP), preserves more than 2500 accessions from over 10 collections performed between 1977 and 1994. As in the rest of the world, most of them are scarcely used due to the poor knowledge of their characteristics and genetic merit, as well as to the almost complete absence of molecular descriptions (Eyhérabide *et al*., 2005; López *et al*., 2005). In this study, we present the morphological and molecular characterization of 30 accessions (17 landraces) of BAP from NWA, collected at sites between 1120 and 2950 masl. They were sown in a common garden at 2300 masl and characterized for 19 morphological characters and 22 SSR loci to evaluate their agronomic potential, determine their genetic constitution and guide *in situ* and *ex situ* conservation efforts.

## Materials and Methods

### Plant material

We selected a set of 30 accessions corresponding to 17 different maize landraces, which covered a broad altitudinal range (collection sites between 1120 and 2950 masl) and represented the morphological diversity occurring in NWA (Fig 1, S1Table).

**Fig 1.**
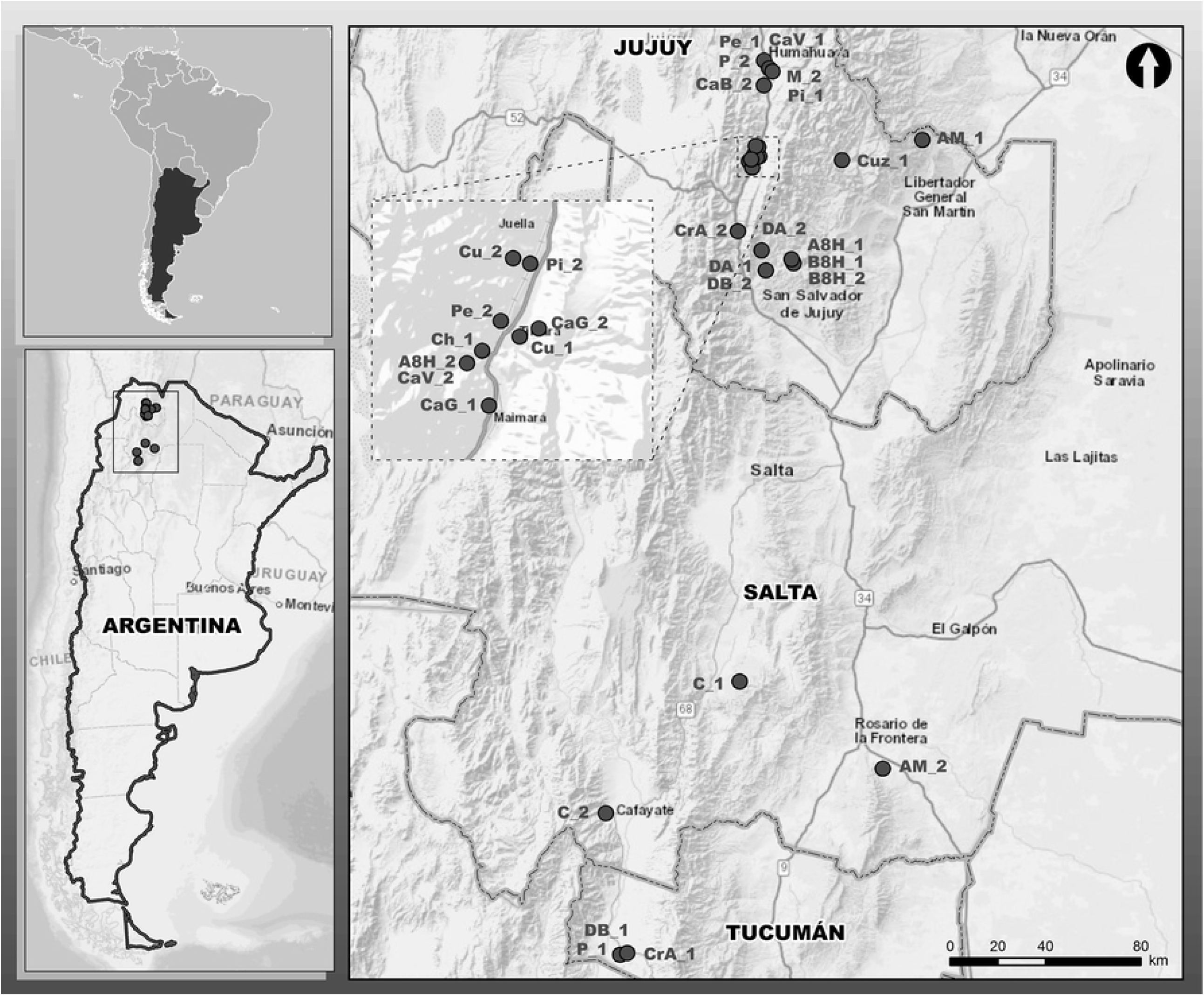
Collection sites of the landraces included in this study. Further details are provided in S1 Table.

### Morphological characterization

All the accessions were morphologically characterized at the Instituto de Pequeña Agricultura Familiar (IPAF), Hornillos, Jujuy Province, Argentina (23°65’17” S, 65°43’55”W; 2300 masl). One-hundred seeds per accession were sown under a randomized block design with two replicates. The elementary plots consisted of two rows of 5 m length, spaced 0.5 m apart. A conventional tillage was applied, with manual weeding and supplementary irrigation. No fertilizer or insecticide was added.

Nineteen agro-morphological descriptors, selected from the list of descriptors of the International Board for Plant Genetic Resources (CIMMYT/IBPGR, Roma, 1991, https://www.bioversityinternational.org/e-library/publications/detail/descriptors-for-maizedescriptores-para-maizdescripteurs-pour-le-mais/), were scored. The following vegetative traits were assessed: plant height (PH), number of leaves (NL), number of leaves above the uppermost ear (NLA), uppermost leaf length (ULL), uppermost leaf width (ULW), venation index (VI) and tillering index (TI). The descriptors measured on the tassel were tassel length (TL), tassel peduncle length (TPL), tassel branching space (TBS), number of primary branches on tassel (NPBT), number of secondary branches on tassel (NSBT) and number of tertiary branches on tassel (NTBT). The variables recorded on the ear were uppermost ear height (EH), ear peduncle length (EPL), ear diameter (ED), mean number of ears per accession (MNE), number of rows of kernels per ear (NRK), and number of kernels per row (NKR).

Vegetative and tassel descriptors were measured on seven to 10 plants per accession per block. The ear and kernel descriptors were measured on 10 ears per accession, which were harvested from the same plants used to evaluate vegetative and tassel descriptors.

### Morphological data analysis

Summary statistics, coefficients of variation (CV) and correlations between traits were computed using InfoStat 2018 [19].

Using the accessions as OTUs, a Principal Component Analysis (PCA) was performed based on average trait values. To avoid biases related to the difference in scale between the variables, data were standardized so that their average was zero and the standard deviation was equal to one. R packages *FactoMineR* [20] and *factoextra* [21] were used to compute and visualize PCA results. Groups of accessions were identified by the K-means algorithm [22], employing the Bayesian Information Criterion (BIC) [23] to find the most likely number of clusters. These analyses were conducted with the R package *adegenet* [24].

### Microsatellite typing

Genomic DNA was extracted from lyophilized young leaves (2–3 days old), germinated from individual kernels following Dellaporta *et al*. [25]. The quality and concentration of the genomic DNA were assessed using NanodropND1000 3.3 software (NanoDrop Technologies^®^).

Twenty-two SSR loci were selected from a preliminary survey of 27, and only loci with unambiguous interpretation were used for this analysis (S2 Table). Genotyping of the SSRs was performed using PCR with fluorescent labeled primers (HEX and FAM). PCR products were size-separated on an Applied Biosystems automated sequencer (ABI 3130 XL) and allele calling was carried out with GeneMapper^®^ 4.0 software (Applied Biosystems, Foster City, USA) using a commercial size standard for allele size assignment (GeneScan ROX 500, Applied Biosystems^®^). Automatic allele calls were subsequently confirmed reviewing all electropherograms.

### Microsatellite data analysis

Mean number of alleles per locus (A), allele frequency, observed (H_o_) and expected (H_e_) heterozygosities, allelic richness (R_s_) [26], presence of population-specific alleles (hereafter referred to as private alleles) and Wright’s fixation indices were assessed using Fstat 2.9.3.2 software [27]. Genetic distances among accessions were computed according to Nei [28] using GeneAlEx 6 (Peakall & Smouse, 2006).

Genetic structure was examined using the Bayesian model-based approach of Pritchard et al. [29] implemented in STRUCTURE 2.3.4 software (http://www.pritch.bsd.uchicago.edu). The number of clusters evaluated ranged from 1 to 10. The analysis was performed using 10 replicates runs per K value, a burn-in period length of 500,000 and a run length of 1,000,000. No prior information on the origin of individuals was used to define the clusters and the correlated frequency model was used for all the analyses. The model assumes that the frequencies in the different populations are likely to be similar due to common ancestry [30]. The deltaK method of Evanno et al [31] was used to identify the most likely number of clusters, using the web tool STRUCTURE HARVESTER [32]. Accessions were assigned to a given cluster when average membership coefficients were higher than an arbitrary cut-off value of 85%. Graphical display of STRUCTURE outputs was performed via the distruct program version 1.1 [33]. Correlations between genetic, morphological, and altitudinal distances were evaluated using Mantel tests (Mantel, 1964). Analyses were conducted based on Spearman correlation coefficients using the *mantel* function of the R package *vegan* [34]. We also inspected Spearman correlations between altitude, morphological traits and genetic diversity measures using InfoStat version 2018 [19]. Significance was assessed with permutation tests (1000 permutations). All graphics were obtained with the R package *ggplot2* [35].

## Results

### Morphological variation

Global averages for the 19 quantitative agro-morphological traits scored in the study are presented in Table 1, along with standard deviations and coefficients of variation. Data for individual accessions are provided in S3 Table.

**Table 1.**
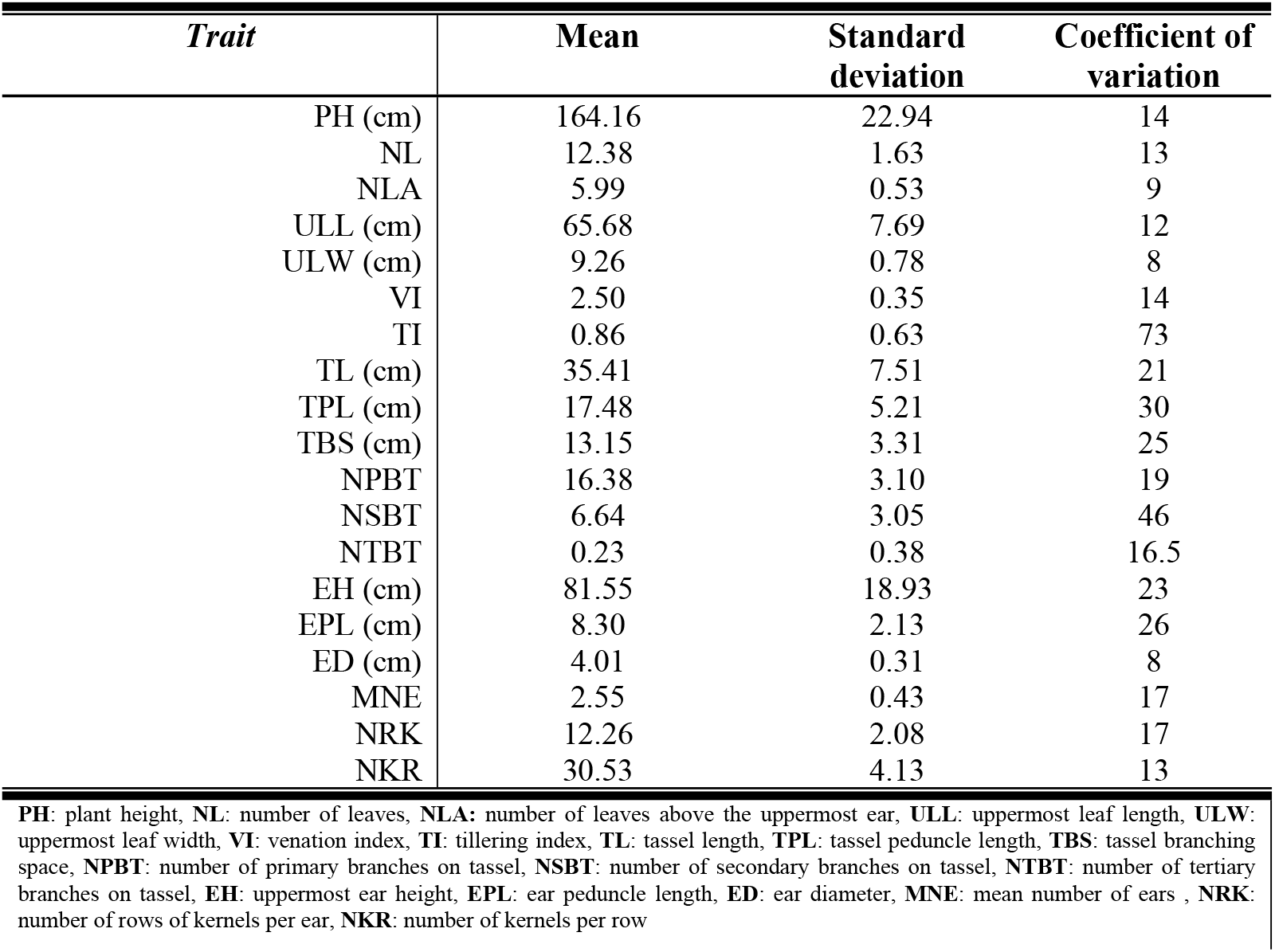
Quantitative trait variation in 30 maize accessions from Northwestern Argentina (NWA).

All the morphological characters analyzed showed considerable levels of variation across accessions (Table 1). The highest coefficients of variation were observed in TI (73%), NSBT (46%) and TPL (30%), while the most homogeneous traits were NLA, ULW and ED, with coefficients of variation of 9, 8 and 8%, respectively.

The analysis of association between variables revealed 56 significant correlations, which decreased to 20 after Bonferroni correction (S4 Table). Most of the significant correlations were moderate, except for PH-EH (r=0.87) and TBS-NSBT (r=0.76).

Analysis of morphological variation using the K-means algorithm allowed the identification of two main groups (K=2, BIC=86.43, diffNgroup criterion) (S3 Table). Accession assignment and distribution along the first two dimensions of the PCA are shown in Fig 2a. The first PC accounted for 32.6% of the variance and was negatively associated with the variables that made the largest contributions to this component, i.e. PH, EH, TBS, and NLA (Fig 2b). The second PC accounted for 13.9% of total variance. It was positively associated with NTBT and negatively correlated with TI, MNE, and ULL.

**Fig 2.**
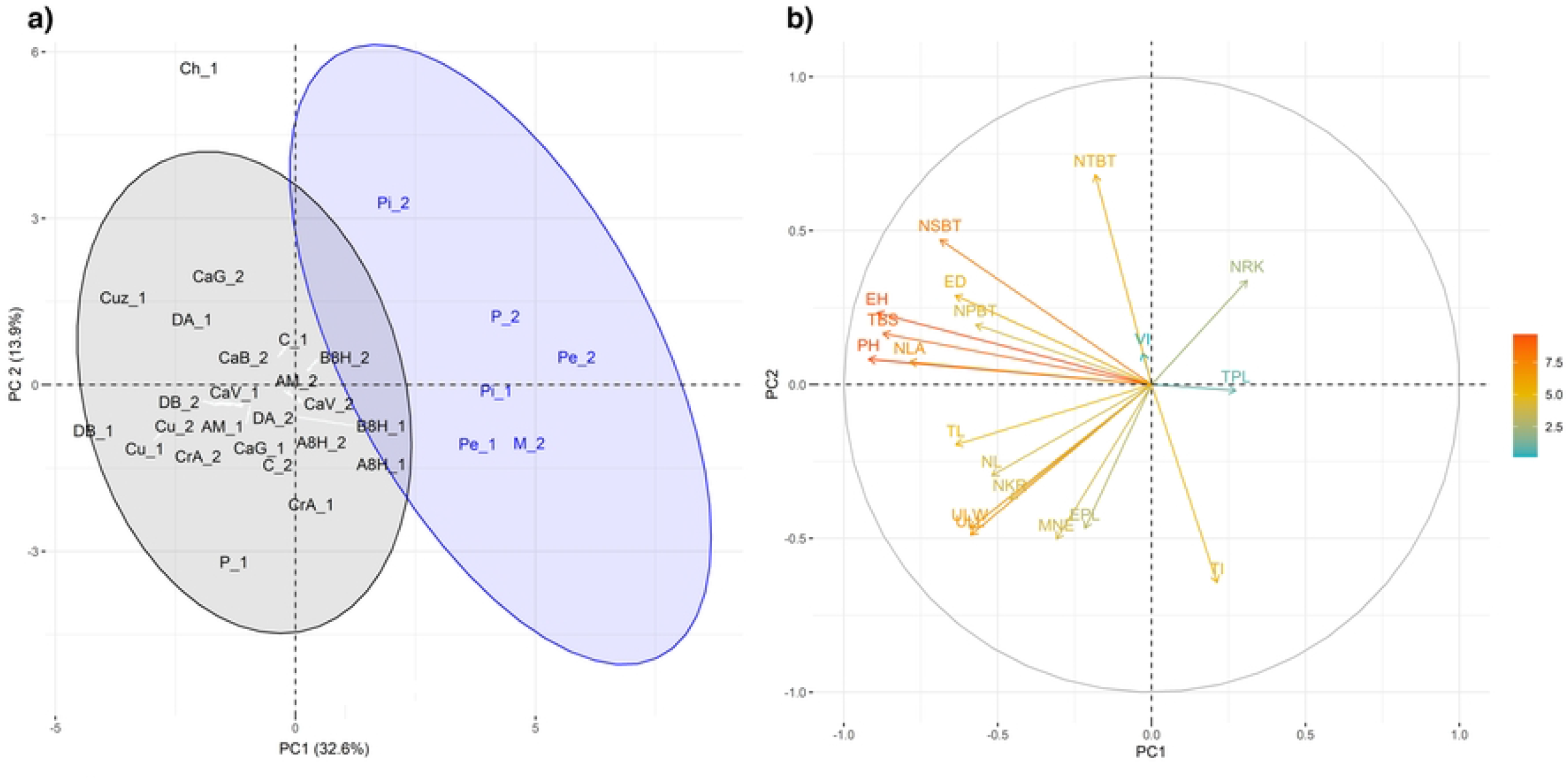
(a) Principal Component Analysis based on 19 agro-morphological traits. Accessions are color-coded according to the groups identified by the K-means procedure. (**b) Correlations and variable contributions to the first two PCs**. The scale corresponds to variable contributions. PH: plant height, NL: number of leaves, NLA: number of leaves above the uppermost ear, ULL: uppermost leaf length, ULW: uppermost leaf width, VI: venation index, TI: tillering index, TL: tassel length, TPL: tassel peduncle length, TBS: tassel branching space, NPBT: number of primary branches on tassel, NSBT: number of secondary branches on tassel, NTBT: number of tertiary branches on tassel, EH: uppermost ear height, EPL: ear peduncle length, ED: ear diameter, MNE: mean number of ears, NRK: number of rows of kernels per ear, NKR: number of kernels per row.

The first group, G1 (black), presented a heterogeneous grain conformation, with accessions showing dent, semi-dent, sweet, flint and floury endosperm types (S1 Table). In this cluster, plants were taller and leafier than those in the second group, G2 (blue). In addition, tassel peduncles were shorter, the tillering index was lower, ear diameter was larger and the average number of grains per ear was 372 (S3 Table).

On the other hand, the G2 group comprised low-height accessions with pop, flint or semi-flint endosperm, fewer leaves, longer tassel peduncles, shorter branching space and very few tassel secondary branches. These accessions produced only a few ears per plant. The ears had smaller mean diameter and fewer grains per row than those in group G1, but a larger number of rows of kernels, resulting in a mean number of grains per ear of 350 (S3 Table).

The third PC (12.1% of total variance) maintained the same groupings, but further divided G1 into two sub-clusters, separating accessions with higher TPL and lower VI from the rest (S1 Fig 1, S5 Table).

### Molecular diversity

Genotyping of 22 SSR loci in 379 individuals revealed a total of 419 alleles (S6 Table). Diversity estimates per locus and accession are provided in S7 Table. The overall number of alleles per locus varied from eight (phi072) to 42 (bnlg244), with a mean of 19.05. The mean number of alleles per locus within accessions was 6.51, ranging from 4.23 (Pi_1) to 8.86 (B8H_2) (Table 2). The index H_o_ varied between 0.52 (CaG_1) and 0.65 (Pi_2), H_e_ between 0.63 (Pi_1) and 0.80 (B8H_2), and Rs between 3.77 (Pi_1) and 5.36 (B8H_2) (Table 2). A total of 35 private alleles were found distributed in 73.3% of the accessions (Table 2), with only four of them showing frequencies higher than 0.1. In regard to He, A and Rs, the most variable accession was B8H_2, while AM_1, Pe_2, DB_1 and Pi_2 exhibited the highest number of private alleles (Table 2). All accessions showed deviations from panmixia, with an excess of homozygotes (Table 2).

**Table 2.** Genetic variability in maize landraces from Northwestern Argentina (22 SSR loci)

| Accession | N | A |  | Ho |  | He |  | Rs |  | PA | F<sub>IS</sub> |
| --- | --- | --- | --- | --- | --- | --- | --- | --- | --- | --- | --- |
|  |  | M | s.d. | M | s.d. | M | s.d. | M | s.d. |  |  |
| AM_1 | 14 | 5.59 | 3.54 | 0.57 | 0.32 | 0.66 | 0.24 | 3.99 | 1.74 | 4 | 0.14** |
| AM_2 | 13 | 5.91 | 3.28 | 0.64 | 0.22 | 0.70 | 0.17 | 4.22 | 1.46 | 1 | 0.09* |
| C_1 | 8 | 5.27 | 2.62 | 0.59 | 0.20 | 0.72 | 0.17 | 4.39 | 1.79 | 0 | 0.18** |
| C_2 | 14 | 6.77 | 2.89 | 0.60 | 0.23 | 0.75 | 0.12 | 4.56 | 1.32 | 0 | 0.19** |
| P_1 | 13 | 8.14 | 3.54 | 0.64 | 0.18 | 0.79 | 0.12 | 5.29 | 1.55 | 2 | 0.20** |
| P_2 | 11 | 5.41 | 3.29 | 0.63 | 0.28 | 0.71 | 0.21 | 4.23 | 1.78 | 0 | 0.11** |
| Pe_1 | 7 | 4.82 | 2.24 | 0.52 | 0.27 | 0.67 | 0.22 | 4.22 | 1.73 | 0 | 0.23* |
| Pe_2 | 15 | 6.32 | 3.67 | 0.55 | 0.23 | 0.67 | 0.24 | 4.21 | 1.77 | 3 | 0.17** |
| DB_1 | 12 | 7.41 | 2.79 | 0.58 | 0.18 | 0.79 | 0.17 | 5.21 | 1.43 | 3 | 0.27** |
| DB_2 | 14 | 7.27 | 3.28 | 0.62 | 0.18 | 0.77 | 0.12 | 4.90 | 1.54 | 1 | 0.20** |
| A8H_1 | 14 | 7.45 | 3.25 | 0.63 | 0.19 | 0.76 | 0.15 | 4.90 | 1.50 | 1 | 0.18** |
| A8H_2 | 15 | 7.55 | 4.42 | 0.62 | 0.22 | 0.72 | 0.23 | 4.77 | 1.96 | 1 | 0.13** |
| CaB_2 | 15 | 6.36 | 3.22 | 0.60 | 0.24 | 0.67 | 0.24 | 4.26 | 1.64 | 1 | 0.10** |
| DA_1 | 15 | 7.36 | 3.06 | 0.61 | 0.24 | 0.77 | 0.13 | 4.92 | 1.39 | 2 | 0.21** |
| DA_2 | 15 | 7.86 | 4.60 | 0.59 | 0.23 | 0.70 | 0.21 | 4.66 | 1.91 | 1 | 0.16** |
| CaV_1 | 12 | 6.50 | 3.69 | 0.55 | 0.25 | 0.69 | 0.25 | 4.54 | 1.91 | 0 | 0.21** |
| CaV_2 | 14 | 6.77 | 3.58 | 0.59 | 0.23 | 0.70 | 0.24 | 4.60 | 1.86 | 2 | 0.15** |
| CrA_1 | 7 | 5.52 | 2.02 | 0.54 | 0.19 | 0.79 | 0.15 | 4.88 | 1.50 | 1 | 0.31** |
| CrA_2 | 15 | 7.23 | 4.39 | 0.63 | 0.23 | 0.69 | 0.24 | 4.53 | 1.88 | 1 | 0.09** |
| B8H_1 | 14 | 7.00 | 3.19 | 0.61 | 0.20 | 0.75 | 0.13 | 4.69 | 1.42 | 0 | 0.18** |
| B8H_2 | 17 | **8.86** | 3.67 | 0.62 | 0.18 | **0.80** | 0.14 | **5.36** | 1.47 | 1 | 0.23** |
| M_2 | 13 | 5.86 | 3.08 | 0.58 | 0.26 | 0.66 | 0.25 | 4.11 | 1.70 | 0 | 0.10** |
| CaG_1 | 8 | 5.59 | 2.97 | 0.52 | 0.25 | 0.70 | 0.28 | 4.64 | 2.01 | 1 | 0.27** |
| CaG_2 | 16 | 6.86 | 3.51 | 0.58 | 0.26 | 0.67 | 0.25 | 4.32 | 1.72 | 1 | 0.13** |
| Pi_1 | 7 | 4.23 | 2.49 | 0.53 | 0.30 | 0.63 | 0.24 | 3.77 | 1.97 | 0 | 0.12* |
| Pi_2 | 14 | 6.36 | 3.57 | **0.65** | 0.23 | 0.72 | 0.20 | 4.45 | 1.61 | 3 | 0.10** |
| Cuz_1 | 11 | 5.95 | 3.53 | 0.53 | 0.27 | 0.65 | 0.27 | 4.33 | 2.06 | 1 | 0.18** |
| Cu_1 | 10 | 5.82 | 3.23 | 0.58 | 0.19 | 0.69 | 0.23 | 4.41 | 1.87 | 2 | 0.16** |
| Cu_2 | 15 | 6.86 | 3.60 | 0.62 | 0.23 | 0.70 | 0.20 | 4.44 | 1.64 | 1 | 0.11** |
| Ch_1 | 11 | 6.38 | 3.35 | 0.57 | 0.27 | 0.71 | 0.22 | 4.55 | 1.76 | 1 | 0.19** |

N: number of individuals. A: number of alleles, Ho: observed heterozygosity, He: expected heterozygosity, Rs: mean allelic richness. PA: private alleles. M: mean, s.d.: standard deviation. The highest value of each index is highlighted in bold. \* p<0.05; \*\* p<0.001.

### Population structure

STRUCTURE-model-based approach allowed discrimination of different genetic clusters within the group of accessions examined here (Fig 3). Using the method of Evanno et al. (2005), maximal ΔK occurred at K= 2, with the next largest peak at K= 3 (S8 Table).

**Fig 3.**
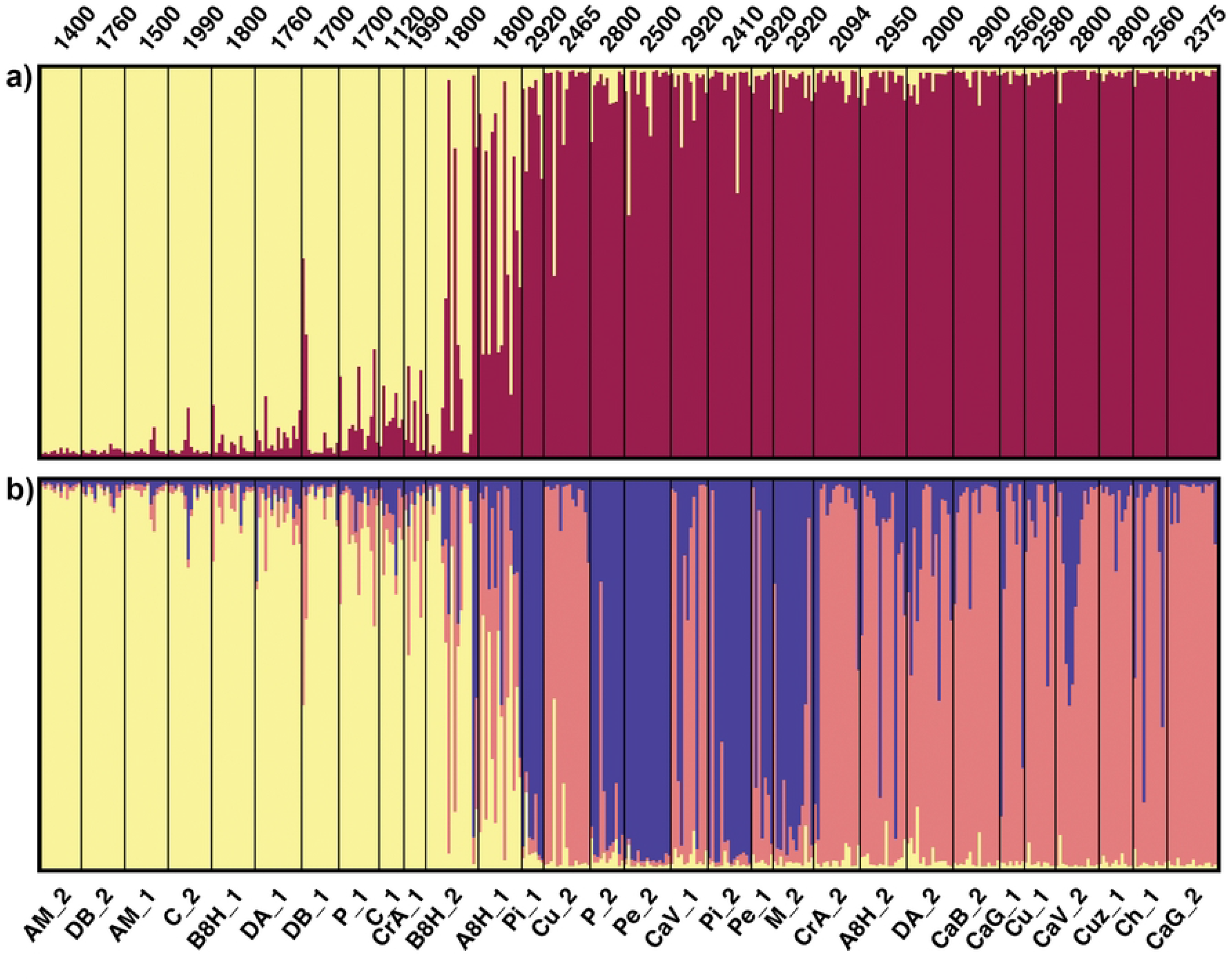
Population structure of maize accessions from Northwestern Argentina (NWA). (**a) STRUCTURE bar plot for K=2. (b) STRUCTURE bar plot for K=3**. Each vertical line represents an individual and colors represent their inferred ancestry from K ancestral populations. Numbers indicate altitude of collection site for each accession (masl).

For K=2, most accessions could be assigned to one of the two groups, with an average proportion of membership higher than 0.85, except for A8H_1 and B8H_2, which showed an admixed composition (Fig 3a), consistent with the high variability indices observed for these accessions. The genetic groups did not match with those obtained with the morphological analysis. The first cluster, yellow, included 10 of the 24 accessions assigned to group G1, while the second cluster, dark purple, included all the accessions of the morphological group G2 and 12 of the group G1 (Fig 3a). When considering K=3, the accessions belonging to the yellow cluster of K=2 still formed a unit, while those of the dark purple cluster became separated into two clusters, blue and pink, one of which coincided with the morphological group G2 (Fig 3b).

A detailed examination of Figure 3a revealed that the genetic groups were associated with the altitude of the collection site, with a clear distinction between accessions cultivated below and above 2000 masl (Fig 3a). Figure 4a shows the proportions of membership to the dark purple cluster (K=2) in function of the altitude of the collection site. Although the correlation was significant (r_Spearman_ = 0.7, p <0.001), the pattern was not gradual, exhibiting an abrupt jump above 2000 masl. When these data were overlapped with the assignments found for K=3, the accessions belonging to the yellow group corresponded to the lowest part of the altitude gradient, while those of the pink and blue clusters were cultivated above 2000 masl. Indeed, for K=3, the mean altitude of the collection site of the accessions within the yellow, pink and blue clusters were 1684, 2584 and 2745 masl, respectively (Fig 4b).

**Fig 4.**
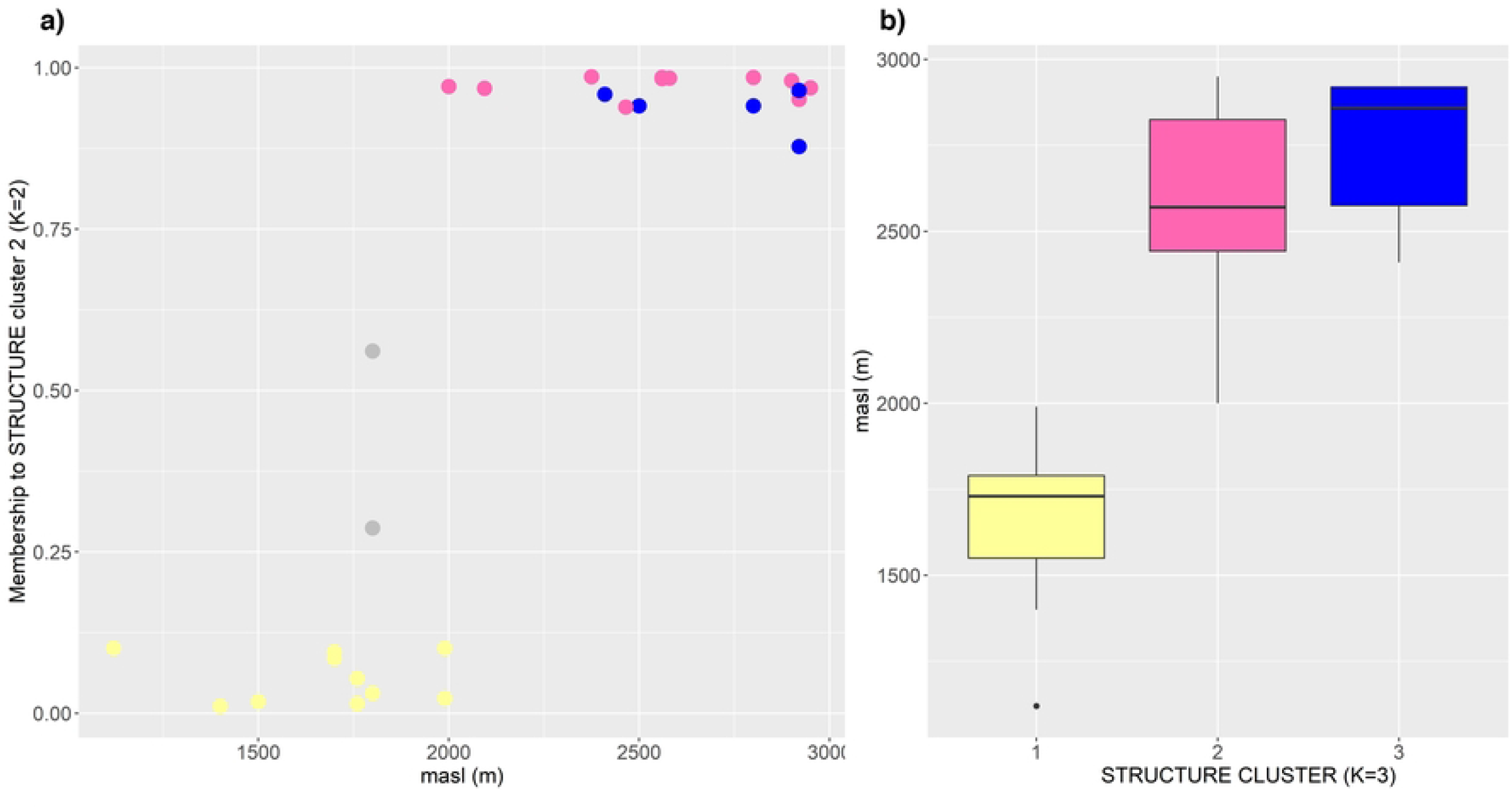
Relationship between population structure and altitude of the collection site. (a) Scatterplot of altitude vs. the inferred ancestry to STRUCTURE cluster 2 (K=2). (b) Box plots of altitude of collection site for the groups inferred with K=3. Accessions are colour-coded according to STRUCTURE assignment for K=3. Accessions in grey could not be assigned to any of the clusters (admixed).

The neighbor-joining network based on Nei’s distances was consistent with the partition into two groups inferred by STRUCTURE (S2 Fig). In most cases, at the level of individual accessions, no clustering was observed between accessions of a same landrace. The overall difference between accessions was moderate, with F_ST_=0.092 (p <0.01).

### Isolation by distance and clinal variation

Morphological distances showed no correlation with genetic, geographic or altitudinal distances among accessions. (Fig 5; Mantel test, p>0.05). Conversely, there were significant associations between genetic and altitudinal distances (Fig 5a), as well as between genetic and geographic distances (Fig 5b) (Mantel test, r=0.44, p<0.001: r=0.54, p<0.001 respectively).

**Fig 5.**
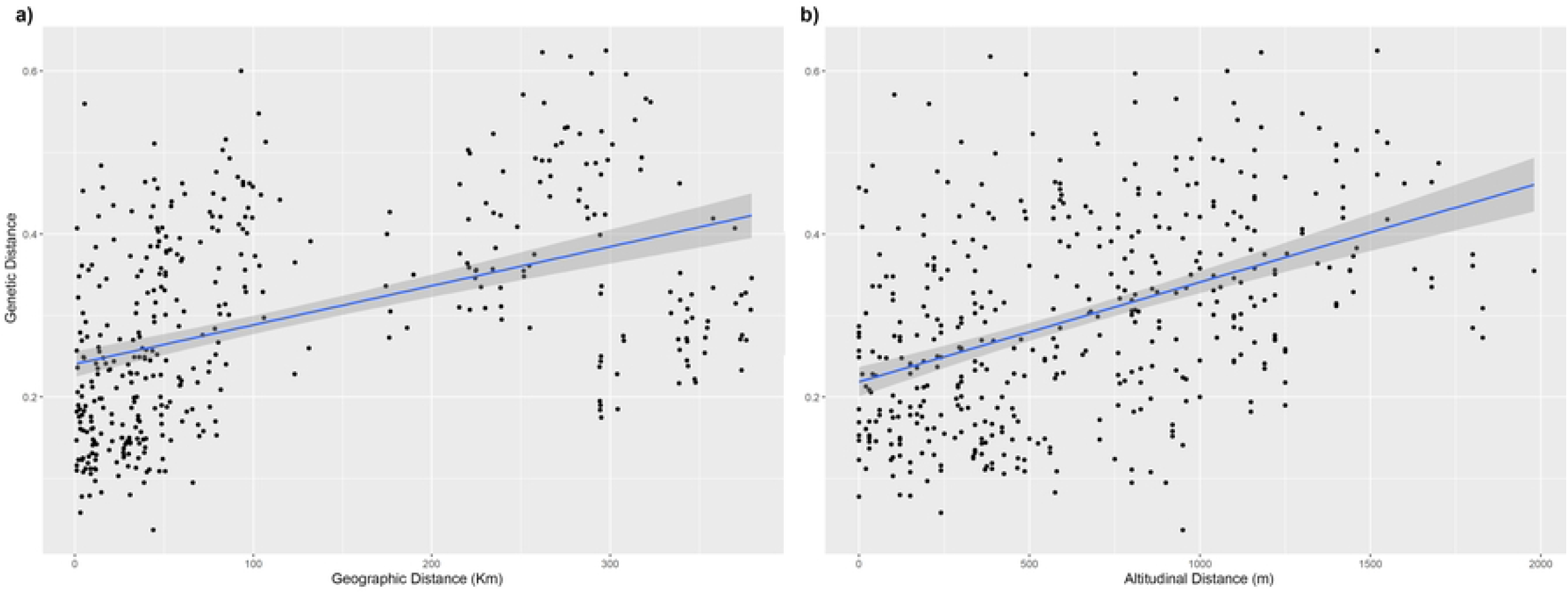
Mantel matrix correlation tests.(a) Scatterplot showing the correlation between genetic and altitudinal distances. (b) Scatterplot showing the correlation between genetic and geographic distances.

Despite the absence of correlation between morphological distance and altitude, the characters NL and NKR were negatively associated with the latter (Table 3). Likewise, a decrease in genetic variability was observed with altitude for the indices He and Rs (Figs 6a and 6b, respectively), but not for the estimators A and PA (Spearman correlation, p>0.05).

**Table 3.** Spearman correlations between agro-morphological traits and altitude.

| Trait | Spearman r | p-value |
| --- | --- | --- |
| PH | -0.28 | 0.1394 |
| EH | -0.32 | 0.0865 |
| NLA | -0.37 | 0.0443 |
| TI | -0.12 | 0.5269 |
| NL | -0.61 | **0.0004** |
| ULL | -0.09 | 0.62 |
| ULW | -0.02 | 0.9017 |
| VI | -0.41 | 0.0229 |
| TL | -0.24 | 0.2014 |
| TPL | 0.39 | 0.0391 |
| TBS | -0.26 | 0.1661 |
| NPBT | -0.48 | 0.0087 |
| NSBT | -0.11 | 0.5875 |
| NTBT | 0.4 | 0.027 |
| EPL | -0.05 | 0.8084 |
| MNE | 1.20E-03 | 0.9948 |
| NRK | 0.06 | 0.7592 |
| ED | -0.07 | 0.7072 |
| NKR | -0.76 | **<0.0001** |

**Fig 6.**
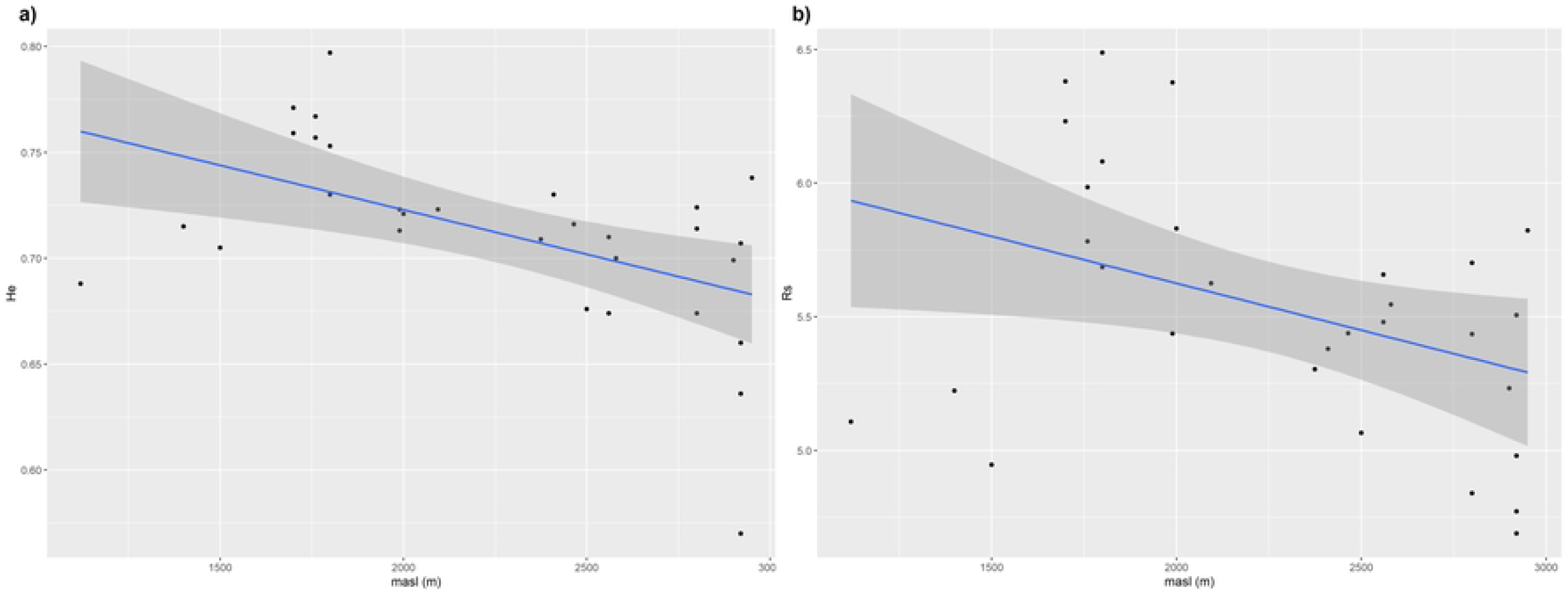
Correlation analysis between diversity estimates and altitude of the collection site. (a) Scatterplot of the correlation between altitude and expected Heterozygosity, He. (b) Scatterplot of the correlation between altitude and allelic richness, Rs.

## Discussion

Efficient use and conservation of the landraces available in germplasm banks requires comprehensive genetic and agronomic characterization focused primarily on variation at the level of individual accessions [3,36–38]. Consistent with this idea, our results suggest that racial assignment, even when performed according to the standardized criteria of germplasm banks, is a poor indicator of similarity between accessions, at both genetic and morphological levels. The lack of racial cohesion may be explained by the dynamics of production and exchange of landraces among local farmers. Small holders in NWA, just like in Mexico [39,40], and most probably in the rest of Latin America, have adopted cultivation practices that allow unintentional racial crossing, as they grow different maize forms or varieties in small areas close to each other, thus facilitating pollen exchange. At the same time, producers usually exchange grains, promoting gene flow through the dispersal of different maize forms among geographically distant localities. Moreover, the introduction of commercial or improved germplasm, which generally provides economic benefits for farmers, has also contributed to blur racial boundaries. In fact, the mixing of such germplasm with native landraces produces creolized varieties that farmers would later identify as “local”.

In this study, the clusters derived from morphological characters are consistent with those previously described for NWA landraces. Melchiorre et al. [41,42] identified two major maize groups: the first one, regarded as more primitive, is related to the popcorn type (Pisingallo) with early maturity, rather corneous grains and smaller plants; the second group is more heterogeneous, with totally or partially corneous or floury grains and larger plants. The landraces studied here display the same pattern, where groups are associated with the type of endosperm and the main discriminating characters are related to plant size (PH, EH, TBS and NLA).

The two major groups identified in the molecular analysis did not coincide with the morphological groups and showed a clear association with the altitude of collection sites, with the morphological group G2 being completely included in the highland group (>2000 masl). The levels of variability also appeared to go along with the differentiation of genetic groups, with a decrease in parameters such as expected heterozygosity and allelic richness in function of altitude. The lower variability of the highland group is consistent with the difficulties in growing maize under harsh environmental conditions. In the highlands, frequent frosts and low temperatures shorten flowering and maturation times, thus reducing seed yields. Consequently, population sizes tend to be smaller, intensifying the effect of genetic drift and promoting the loss of diversity.

Regardless of how variability is distributed, NWA landraces taken as a whole show high levels of diversity, which are similar to the values recorded at the centers of crop origin. The SSR loci analyzed here are a subset of those evaluated by Vigouroux et al. [5]. When limiting the comparisons to this subset of loci, the mean number of alleles detected in the present study (A=19.05) is slightly lower than that reported by Vigouroux et al. (2008) for both the Highland Mexico (A=20.4) and Andean (A=21.45) groups. Likewise, the estimates of He varied within a range previously reported for landraces from all over the Americas (0.61-0.81) [5,9,10,39,43,44]. These results emphasize the potential value of the NWA landraces conserved in the BAP, since only 30 accessions contained almost the same number of alleles as germplasm coming from the central regions of maize distribution.

Several studies have found a relationship between altitude of collection site and genetic composition of maize landraces both at the regional and continental levels [8,10,45]. The existence of correlation between genetic differentiation and altitude may be interpreted as resulting from clinal variation, where the genetic distances increase gradually with increasing altitudinal distance. However, in the landraces analyzed here, the relationship between altitude and inferred ancestry to the clusters obtained by STRUCTURE (Fig 4a) suggests a more abrupt cutoff, where the accessions of intermediate position are those showing a signature of recent introgression.

On the other hand, although morphological differentiation was not found to be correlated with altitude, individual trait analysis revealed a significant decrease for NL and NRK. Considering that all measurements were made in a common garden experiment at 2300 masl, these results may be interpreted as a local adaptation rather than phenotypic plasticity in response to environmental conditions. In fact, the relationship between flowering time, which is a key factor in the adaptation of landraces to altitude [46,47], and NL has been extensively documented [48–52]. In particular, Li et al [53] showed that flowering time shared genetic determinants with the number of leaves below, but not above, the uppermost ear (NLA). These results may help explain why we found an association between NL, but not NLA, and altitude.

As observed here for NL, the reduction in genome size with altitude has been proposed to be an indirect consequence of selection on flowering time [54]. In agreement with these hypotheses, NWA landraces also exhibit patterns of clinal variation for both the DNA content of autosomes and the occurrence of B chromosomes [17,55]. Evidence of their adaptive significance was provided by Lia et al. (2007).

Alternative scenarios can be postulated to explain the genetic structuring pattern found in this study, i.e., the existence of two well-differentiated groups, one below and the other above 2000 masl, in a relatively small area. Rivas [56] compared the genetic relationships between the materials analyzed here and those included in the studies of Vigouroux et al. [5], Lia et al. [16] and Bracco et al. [57]. This author found that the landraces at the lower part of the altitudinal gradient were assigned to the Tropical Lowland cluster [5], while those from the highland zone were assigned to the Andean cluster. These two groups have been related to distinct events of maize dispersal in South America [58,59], suggesting the current coexistence of two genetic pools of different age in NWA, namely, landraces that entered through the eastern lowlands of the continent and those introduced from the Central Andes. Indeed, Vigouroux et al. [5] had already considered the idea of a secondary contact zone encompassing Bolivia, Argentina, Paraguay and Uruguay, where typical Andean landraces became mixed with those from the eastern lowlands of South America. More recently, Kistler et al. [60] proposed the existence of three major maize lineages in South America. Two of these, the Andean and Lowland lineages, derived from a partially domesticated maize that had been introduced to south-western Amazon at early stages. The third lineage, Pan-American, was introduced later through lowland South America. In this scenario, available information is still insufficient to establish putative relationships between the lowland lineages postulated by Kistler et al. [60] and the lowland NWA landraces. In addition to the uncertainties concerning the movements of maize during pre-Columbian times, the influence of the varieties and hybrids resulting from modern breeding is another factor to be considered when interpreting the genetic composition of maize from NWA. In this regard, Cámara Hernández et al. [13] provided numerous examples of creolized landraces, particularly for the lowlands. Likewise, evidence obtained with chloroplast genome sequencing support the occurrence of introgression between traditional landraces and more modernly improved varieties [61].

In conclusion, the present study constitutes a valuable contribution to existing *ex situ* conservation programs and to design future collection and management strategies. In particular, our results indicate that altitudinal structuring is a key factor in decision-making. For example, Perales et al. [62] reported that the replacement rate of native landraces by modern varieties in Mexico was considerably higher in lowlands and mid-elevation sites than in highlands because there were few modern varieties capable of outperforming native materials in high-altitude environments. These findings led them to suggest that interventions should favor *in situ* conservation strategies in lowland or intermediate elevation sites, where modern varieties have largely outcompeted native landraces. Despite the lack of similar studies in NWA landraces, it is clear that the altitudinal structuring of genetic diversity constitutes a relevant factor to be considered for their conservation.

## Acknowledgments

We thank all the collaborators at the Instituto de Pequeña Agricultura Familiar (IPAF) for their field assistance. We are also grateful to Dr. Silvia Pietrokovsky, who kindly revised the English of the manuscript.

## Supporting information

S1 Fig. This is the S1 Fig Title. This is the S1 Fig legend. S2 Fig. This is the S2 Fig Title. This is the S2 Fig legend.

S1 Table. This is the S1 Table Title. This is the S1 Table legend. Fig_S1_PCA_individuos_PC3_PC1

Fig_S2_NJ_sssr_379.tree

1_Supplementary Table_S1_Accessions

2_Supplementary Table_S2_primers

3_Supplementary Table S3_Morphologicaldata

4_Supplementary Table_S4_Correlations

5_Supplementary Table_S5_PCA_contributions

6_Supplementary Table_S6_SSR_data

7_Supplementary Table_S7_Genetic_diversity

8_Supplementary_Table_S8_Evanno_deltaK

